# Free fatty acid 2 receptor regulates the NADPH oxidase activity induced by formyl peptide receptor specific agonists

**DOI:** 10.64898/2026.09.05.749594

**Authors:** Danni Wang, Lena Björkman, Claes Dahlgren, Huamei Forsman

## Abstract

The neutrophil NADPH oxidase is activated by signals generated by formyl peptide receptor (FPR) agonists recognized by FPR1 (fMLF) and FPR2 (WKYMVM), respectively. Also, the antagonists cyclosporin H (specific for FPR1) and PBP10 (specific for FPR2) inhibit the NADPH oxidase activity when induced by the two peptide agonists. When bound to its receptor, the non-activating positive allosteric modulator Cmp58, specific for the free fatty acid 2 receptor (FFA2R), affects not only the response induced by agonists specific for FFA2R, but also the activating potency but not the efficacy of the two FPR activating peptides. Even though Cmp58 is without effect on the efficacy of the response induced by the FPR agonists, the inhibitory effect of the respective FPR specific antagonists is reduced. This sensitivity shift was reversed by an FFA2R specific antagonist suggesting that two different signals generated by the FPRs activate the NADPH oxidase. According to a receptor trans-regulation model, the activated FPRs generate signals that directly activate the NADPH oxidase and signals that activate the allosterically modulated FFA2Rs to elicit activation of the NADPH.

## Introduction

The closely related formyl peptide receptors (FPRs) belong to a subfamily of G protein coupled receptor (GPCRs) and comprises three human members, FPR1, FPR2, and FPR3 (1, 2). Two of these (FPR1 and FPR2), that are expressed in human neutrophils, have been used as model GPCRs in many studies on the role of different inflammatory agents in innate immune functions and regulation of inflammatory reactions. Endogenous agonists that are recognized by the neutrophil FPRs originate from mitochondrial proteins and are generated during tissue destruction whereas exogenous agonists are generated by invading microbes (3). Several synthetic FPR specific tool compounds have also been identified and characterized over the years (4, 5). The two neutrophil FPRs exhibit a large overall similarity not only in amino acid sequence but also in their sub-cellular localization, signaling properties and in the receptor down-stream functions that they regulate. Neutrophils contain granule/vesicle-localized storage-pools of the FPRs that can be mobilized to the plasma membrane by priming agents such as tumor necrosis factor (TNF), they couple to Gα_i_ containing G proteins that initiate signaling pathways that include a phospholipase C catalyzed rise in the cytosolic concentration of free calcium ions ([Ca^2+^]_i_). Functionally the FPR agonists are commonly used potent secretagogues, chemoattractants and, activators of the neutrophil superoxide anion (O_2_^-^) generating NADPH oxidase system. Two of the commonly used FPR agonists are the N-formylated tripeptide fMet-Leu-Phe (fMLF – FPR1 agonist) and the hexapeptide Trp-Lys-Tyr-Met-Val-Met (WKYMVM – FPR2 agonist) (2, 6). Two potent antagonists specific for FPR1 (cyclosporin H; (7)) and FPR2 (PBP10; (8, 9)), respectively, are tool compounds available for studies of the role of the two FPRs in activation and signaling in human neutrophils.

The Free Fatty Acid 2 Receptor (FFA2R earlier termed GPR43) is also a highly expressed GPCR in human neutrophils (10), but in contrast to the robust effect of FPR agonists, activation of FFA2R by one of its orthosteric agonists, the short-chain fatty acids acetate, propionate, and butyrate, generates only low levels of NADPH oxidase activity (11). The function in neutrophils and in innate immunity/inflammation of FFA2R has not yet been elucidated but the identification of potent tool compounds such as various allosteric modulators including Cmp58 and the specific antagonist CATPB, specific for FFA2R, has opened for basic studies of signaling and function of this receptor (12–14). The basic concept of positive allosteric GPCR modulator is fulfilled by Cmp58; this allosteric modulator binds to a receptor site in FFA2R that is structurally separated from the binding site for orthosteric agonists, and when bound, the responses induced by orthosteric FFA2R agonists are increased (15). The allosteric priming effect is, however, not restricted to ligands recognized by FFA2R. The non-activating allosteric FFA2R modulator Cmp58 has off-target effects on the response induced by several different GPCRs including normally non- as well as low-activating agonist recognized by the P2Y_2_receptor (receptor for ATP), the PAF receptor (receptor for platelet activating factor), the HCA_3_R (receptor for hydroxy-carboxylicacid) and, the BLT1receptor (receptor for leukotriene B4) (16–18). The proposed molecular mechanism by which the allosterically modulated FFA2R affects signaling properties of non-/low-activating GPCRs opens for the hypothesis, that signaling also by other neutrophil GPCRs may be regulated by FFA2R. The recently described transactivation data, showing that the allosterically modulated FFA2Rs are activated by the agonist occupied C5aR (the receptor for the complement component C5a) strongly support this hypothesis (19).

In this study we have investigated effects of the FFA2R specific PAM Cmp58, on the neutrophil NADPH oxidase activity mediated by the two FPR specific peptide agonists fMLF and WKYMVM. In agreement with earlier findings, we show that the FPR agonists alone trigger a strong NADPH oxidase response, responses that were inhibited by the respective FPR1/FPR2 antagonist. The allosteric FFA2R modulator Cmp58 was without effect of the efficacy of the FPR agonist induced response but increased their potency. In addition, we found that the allosteric FFA2R modulator also changed the inhibitory profile of the respective antagonist on the FPR1/FPR2-mediated response, an effect that was reversed by the FFA2R specific antagonist CATPB. These data suggest that the neutrophil FFA2R is a master regulator of GPCR signaling and this is achieved through a novel receptor transactivation mechanism.

## 2. Materials and Methods

### 2.1 Chemicals

Dextran T500 and Ficoll-Paque Plus Medium used for the isolation of neutrophils were obtained from Pharmacosmos (Holbaek, Denmark) and Fischer Scientific (Gothenburg, Sweden), respectively. Isoluminol, TNF, horseradish peroxidase (HRP), compound 58 (Cmp58; (S)-2-(4-chlorophenyl)-3,3-dimethyl-N-(5-phenylthiazol-2-yl)butanamide), fMLF, WKYMVM, cyclosporin H, and PBP10 were purchased from Sigma-Aldrich (Merck, Burlington, MA, USA). CATPB ((S)-3-(2-(3-chlorophenyl)acetamido)-4-(4- (trifluoromethyl)phenyl) butanoic acid) was from Tocris (Bristol, UK). Stock solutions of the peptides were prepared in DMSO, and working solutions were prepared in Krebs-Ringer glucose phosphate buffer (KRG, 120 mM NaCl, 4.9 mM KCl, 1.7 mM KH_2_PO_4_, 8.3 mM Na_2_HPO_4_, 1.5 mM MgSO_4_, 10 mM glucose, and 1 mM CaCl_2_ in dH_2_O, pH 7.3).

### 2.2 Isolation of human neutrophils

Human neutrophils were isolated from buffy coats that were obtained from anonymous healthy human blood donors at the blood bank of Sahlgrenska University Hospital in Gothenburg, Sweden. As earlier described, dextran sedimentation of red blood cells followed by Ficoll-Paque gradient centrifugation was used to isolate neutrophils (20). After separation, contaminating red blood cells were lysed by hypotonic lysis and the remaining cells were washed and resuspended (1 x 10^6^ cells/mL) in KRG and kept on ice until used in further assays on the same day as the isolation. The purity of neutrophils was determined in an automatic analyzer (Sysmex KX-21 N Hematology Analyzer, Sysmex Corporation) and routinely contained ≥ 90% neutrophils. To amplify the activation signals of the neutrophil NADPH oxidase, freshly isolated neutrophils were primed with TNF (10 ng/mL; 1 x 10^6^ cells/mL) at 37°C for 20 minutes in a water bath and then stored on ice until use.

### 2.3 Measuring NADPH oxidase activity in neutrophils

An isoluminol-enhanced chemiluminescence (CL) system was used to measure the production of superoxide anions (O_2_^-^) generated by the neutrophil electron transporting NADPH oxidase as described earlier (21, 22). All the measurements were performed using a six-channel Biolumat LB 9505 (Berthold Co., Wildbad, Germany). Disposable polypropylene tubes (4 mL) were used, containing a 900 µL reaction mixture comprising 1 x 10^5^ neutrophils, isoluminol (2 x 10^-5^ M) and HRP (4 units/mL). The tubes were incubated for 5 min at 37°C before adding the activating ligand (100 µL), and the light emission was recorded continuously. To determine the effects of receptor-specific antagonists and allosteric modulators, these ligands were added to the reaction mixture 1-5 min before agonist stimulation. Controls were run in parallel for comparison. The O_2_^-^ production, recorded continuously over time, was expressed in Mega counts per minute (Mcpm) and peak activities were used for comparisons of the NADPH oxidase activities.

### 2.4 Data analysis

The raw data were processed using GraphPad Prism 10.6 (GraphPad Software, San Diego, CA, USA). The results obtained from the data are presented as mean with standard error of the mean (SEM). The number of repeats performed independently on neutrophils isolated from different individuals (n) for each figure is described in the figure legends.

## Results

### The peptides fMLF and WKYMVM are potent FPR agonists that induce an activation of the NADPH oxidase in TNF primed neutrophils

A superoxide (O_2_^-^) generating electron-transporting NADPH oxidase is expressed in neutrophil phagocytes (23). When activated, this enzyme system transfers electrons from NADPH generated by the hexose monophosphate in the neutrophil cytosol to extracellularly localized molecular oxygen that is reduced to O_2_^-^. Based on previous published data (16–18), showing that TNF priming is of importance in a novel receptor transactivation mechanism involving FFA2R, TNF-primed neutrophils were used in the experiments described below, designed to determine FFA2R-dependent trans-regulating mechanisms in neutrophils activated by peptide agonists recognized by formyl peptide receptor 1 and 2 (FPR1; FPR2), respectively. The two peptides fMLF (FPR1 specific) and WKYMVM (FPR2 specific) are potent agonists that through an interaction with their respective receptor dose-dependently activate the NADPH

### The allosteric FFA2R modulator Cmp58 increases the potency of the FPR activating peptides

Contrary to the general belief regarding the receptor selectivity for allosteric GPCR modulators, the allosteric FFA2R modulator Cmp58 turns not only propionate, a non-activating orthosteric agonist recognized by FFA2R, into an NADPH oxidase activating ligand, but Cmp58 also positively modulated the response induced by several agonists recognized by other neutrophil GPCRs (16). The mechanisms by which FFA2R regulates the function of other GPCRs has not yet been disclosed but a working model for how signals generated by one neutrophil GPCR affect the activity of FFA2R has been presented (see the discussion and (3)). To determine if Cmp58 affects the activation pattern of the NADPH oxidase when induced by different concentrations of the FPR agonists, the allosteric modulator was included in the system designed to measure generation and release of O_2_^-^. The data obtained show that when neutrophils were activated by low concentrations of the FPR1 and FPR2 agonists, Cmp58 increased the neutrophil response (Fig 2). Cmp58 did, however, not affect the magnitude of the response induced by high concentrations of fMLF (Fig 2A) or WKYMVM (Fig 2B), concentrations that on their own, fully activate the NADPH oxidase (Fig 1).

**Figure 1.**
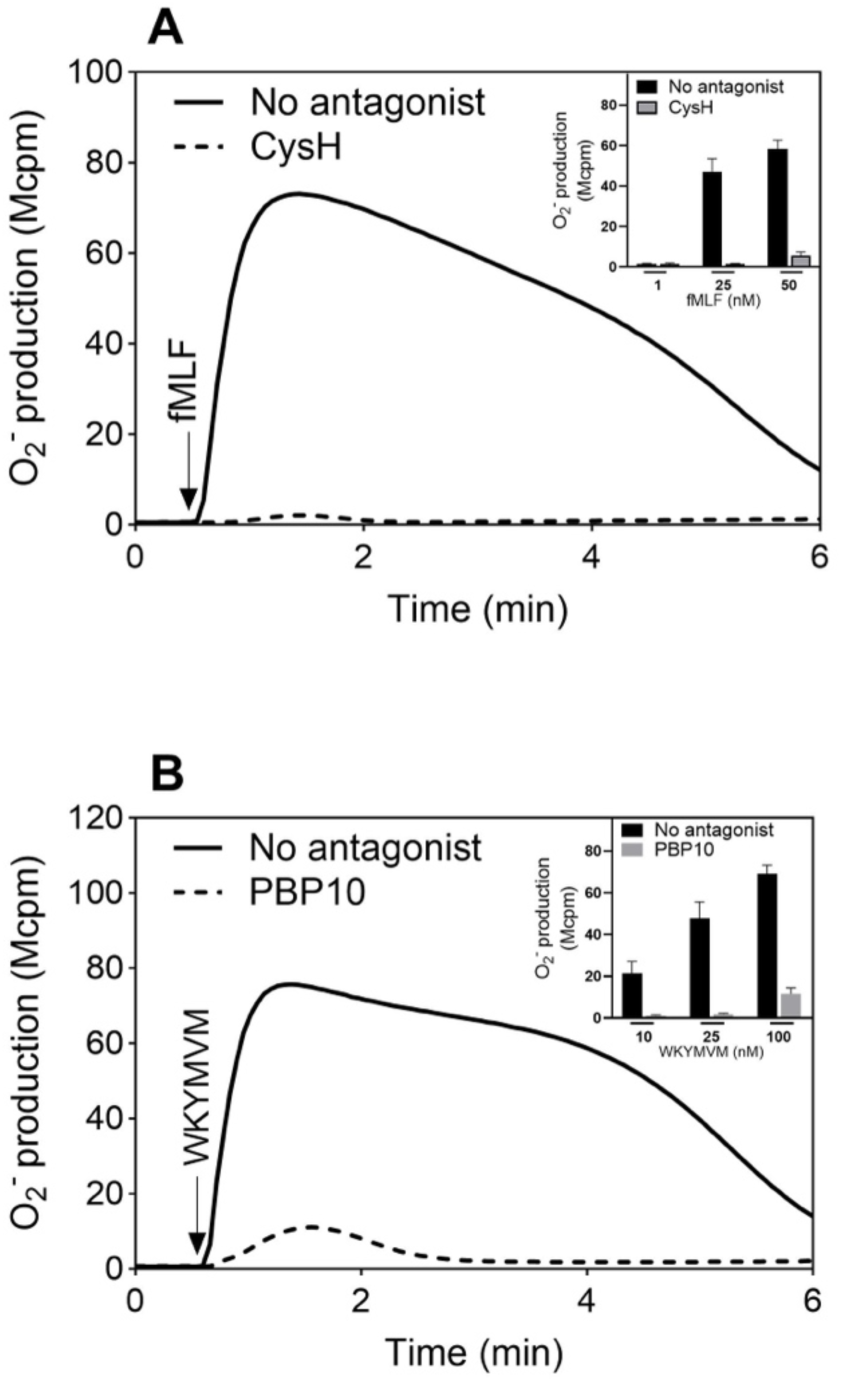
TNF primed neutrophils activated by FPR agonists and inhibition of the response by receptor specific antagonists. **(A)** The neutrophil response induced by the FPR1 agonist fMLF (50 nM), added at the time point marked with an arrow. The response was measured in the absence (solid line) or presence (broken line) of the FPR1 specific antagonist cyclosporin H (CysH; 1µM). **Inset:** The inhibitory effect of CysH (1µM) on the response induced by three different concentrations of the FPR1 agonist fMLF. The NADPH oxidase activity is expressed as the peak values (Mcpm; mean ± SEM; n > 3) of O_2_^-^ production induced by fMLF. **(B)** The neutrophil response induced by the FPR2 agonist WKYMVM (100nM), added at the time point marked with an arrow. The response was measured in the absence (solid line) or presence (broken line) of the FPR2 specific antagonist PBP10 (1µM). **Inset:** The inhibitory effect of PBP10 (1µM) on the response induced by three different concentrations of the FPR2 agonist WKYMVM. The NADPH oxidase activity is expressed as peak values (Mcpm; mean ± SEM; n>3) of O_2_^-^ production induced by WKYMVM.

**Figure 2.**
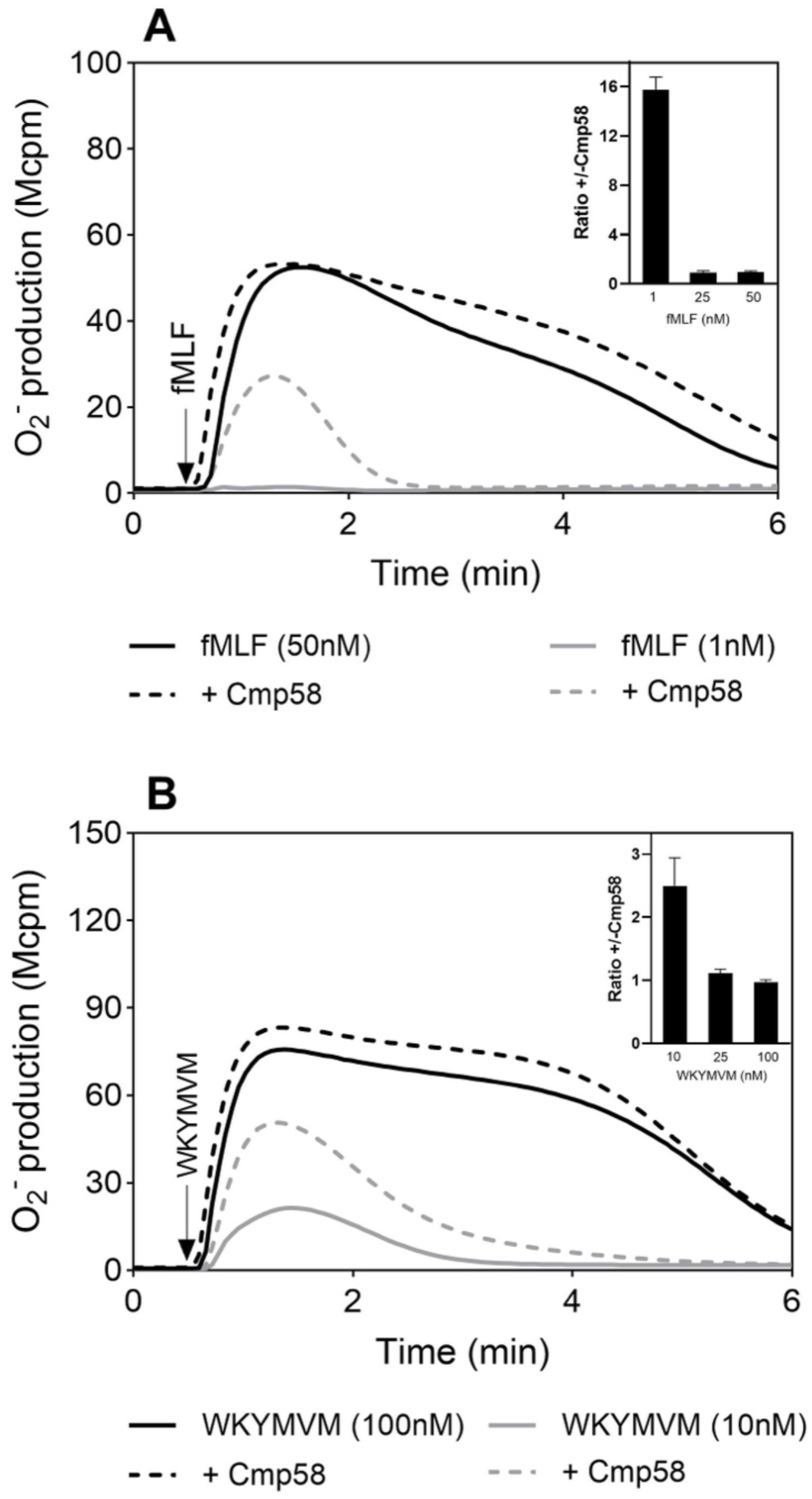
Effect of an FFA2R specific positive allosteric modulator (Cmp58) on the neutrophil response induce by FPR agonists. Neutrophils were incubated without or with Cmp58 (1 µM; 5 min at 37°C) and the O_2_^-^ production was measured continuously following an activation by two different concentrations of an FPR agonist. **(A)** The response induced by fMLF (50nM, black line; 1nM, grey line) was determined in the absence (solid lines) and presence of Cmp58 (broken lines). **Inset:** The effect of Cmp58 (1µM) on the response induced by three different concentrations of fMLF. The NADPH oxidase activity is expressed as the ratio between the peak values of the response in the presence (+) and absence (-) of Cmp58 (mean ± SEM; n=4). **(B)** The response induced by WKYMVM (100nM, black line; 10nM, grey line) was determined in the absence (solid lines) and presence of Cmp58 (broken lines). **Inset:** The effect of Cmp58 (1µM) on the response induced by three different concentrations of WKYMVM. The NADPH oxidase activity is expressed as the ratio between the peak values of the response in the presence (+) and absence (-) of Cmp58 (mean ± SEM; n>4).

### Inhibition by the specific FFA2R antagonist CATPB on the response induced by FPR agonist in the presence of Cmp58

It has been shown that the binding site for the allosteric modulator Cmp58 overlaps with the orthosteric binding site in FFA2R (17) and by that, its binding and allosteric effect is inhibited by CATPB, an FFA2R antagonist that binds to the orthosteric site in the receptor (17). In agreement with the receptor specificity of CATPB, the Cmp58 primed neutrophil response induced by low concentrations of fMLF and WKYMVM was inhibited by CATPB (Fig 3).

**Figure 3.**
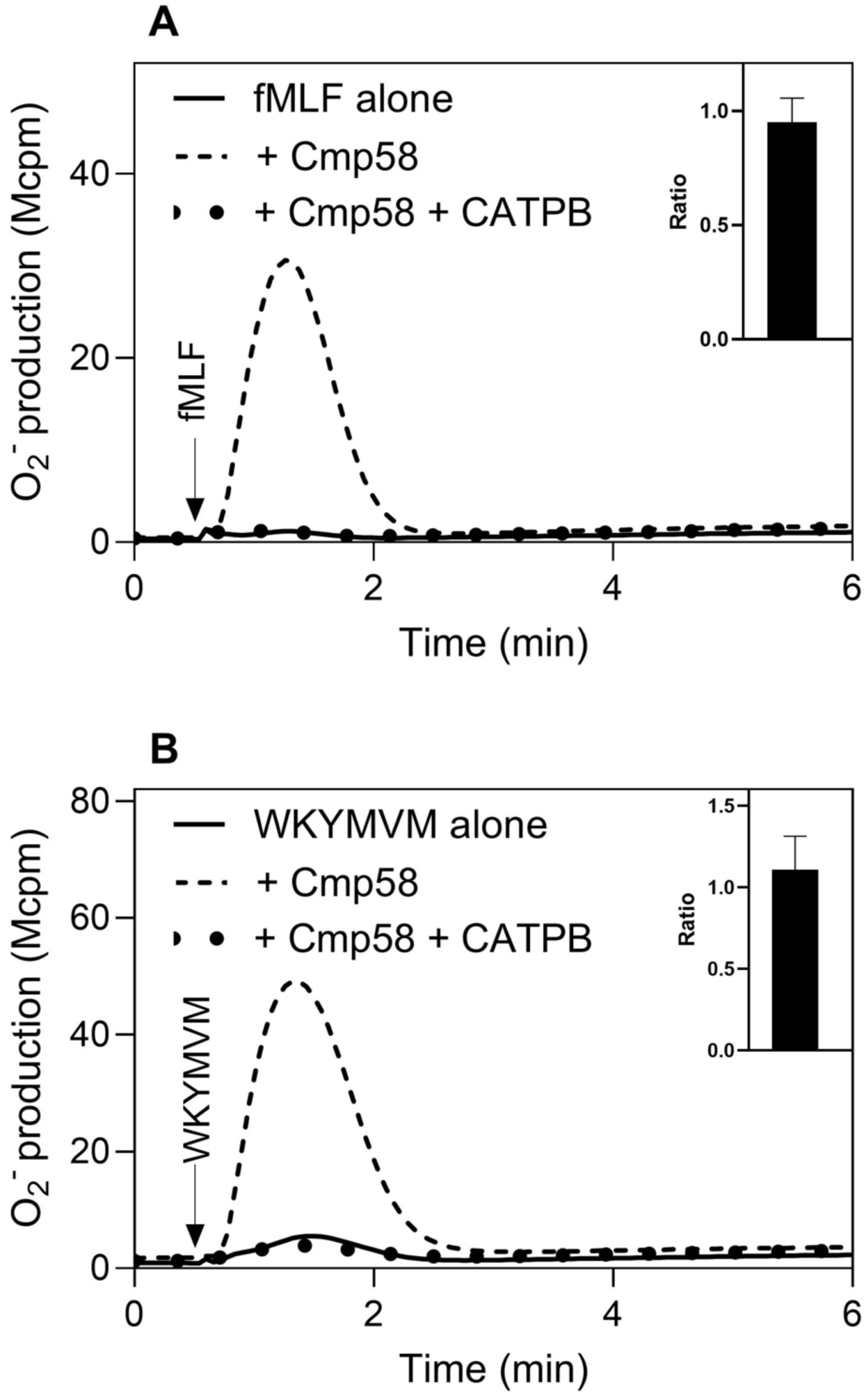
Effects of the FFA2R specific antagonist CATPB on the response induced by low concentrations of FPR agonists in the presence of the positive allosteric FFA2R modulator Cmp58. Neutrophils were incubated without or with Cmp58 (1 µM; 5 min at 37°C), either alone or in combination with the FFA2R antagonist CATPB (100nM), and O_2_^-^ production was measured continuously following an activation by an FPR agonist. (A) The response induced by 1nM fMLF in the absence of Cmp58 (solid line) and in the presence of Cmp58 alone (broken line) or in combination with CATPB (dotted line). Inset: The effect of CATPB (100nM) on the response induced by fMLF expressed as the ratio between the peak values of the response to the agonist alone and those obtained in the presence of both Cmp58 and CATPB (mean ± SEM; n=3). (B) The response induced by 10nM WKYMVM in the absence of Cmp58 (solid line) and in the presence of Cmp58 alone (broken line) or in combination with CATPB (dotted line). Inset: The effect of CATPB (100nM) on the response induced by WKYMVM expressed as the ratio between the peak values of the response to the agonist alone and those obtained in the presence of both Cmp58 and CATPB (mean ± SEM; n=3).

The primed response levels of the activity in the presence CATPB returned to the those in the absence of Cmp58 (Fig 3 insets in A and B). These data validate the two-receptor transactivation model suggested for activation of the allosterically modulated FFA2R, by signals generated by other GPCRs (see the discussion and (3)). In contrast, the response induced by a high concentration of fMLF and WKYMVM, respectively, was not affected by the FFA2R antagonist CATPB (Fig 4).

**Figure 4.**
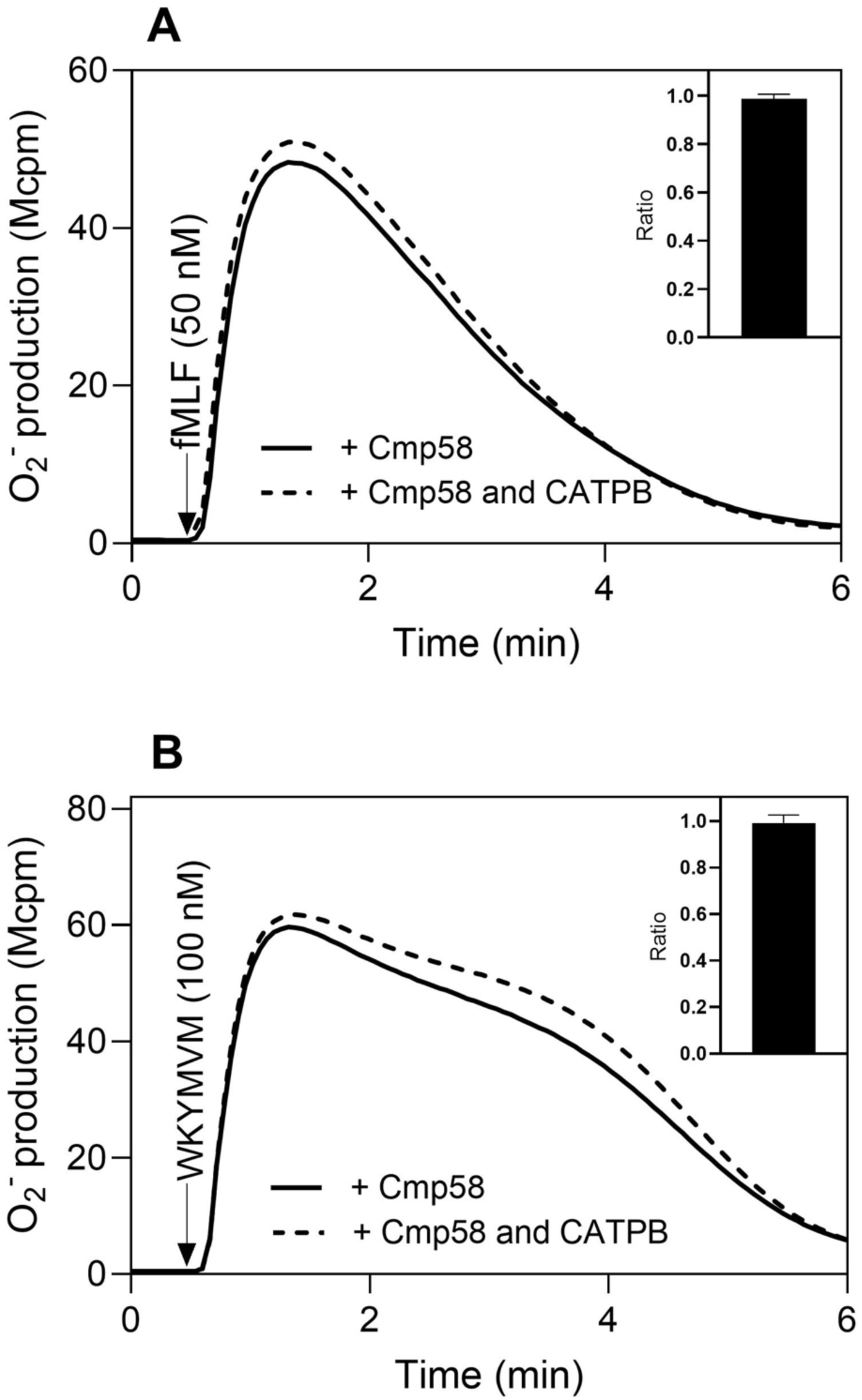
Effects of the FFA2R specific antagonist CATPB on the response induced by high concentrations of FPR agonists in the presence of the positive allosteric FFA2R modulator Cmp58. Neutrophils were incubated with Cmp58 (1 µM; 5 min at 37°C) alone or in combination with the FFA2R antagonist CATPB (100 nM), and the O_2_^-^ production was measured continuously following an activation by an FPR agonist. The response induced by 50 nM fMLF (**A**) or 100 nM WKYMVM (**B**) in the presence of Cmp58 alone (solid lines) or in combination with CATPB (dashed lines) is shown. **Inset:** The effect of CATPB (100 nM) on the response induced by 50 nM fMLF (**A**) or 100 nM WKYMVM (**B**), expressed as the ratio between the peak values of the response in the presence (+) and absence (-) of CATPB (mean ± SEM; n=5)

### Reduced inhibitory effect of the FPR specific antagonists cyclosporin H and PBP10 in the presence of Cmp58

The lack of inhibition mediated by CATPB on the response induced by high concentrations of the FPR agonists (see Fig 4) agree with the results showing that Cmp58 had no effect on the neutrophil response induced by high concentrations of the FPR agonist (see Fig 2). These responses were, however, not fully inhibited by the FPR1 and FPR2 specific antagonists cyclosporin H (Fig 5A) and PBP10 (Fig 5B).

**Figure 5.**
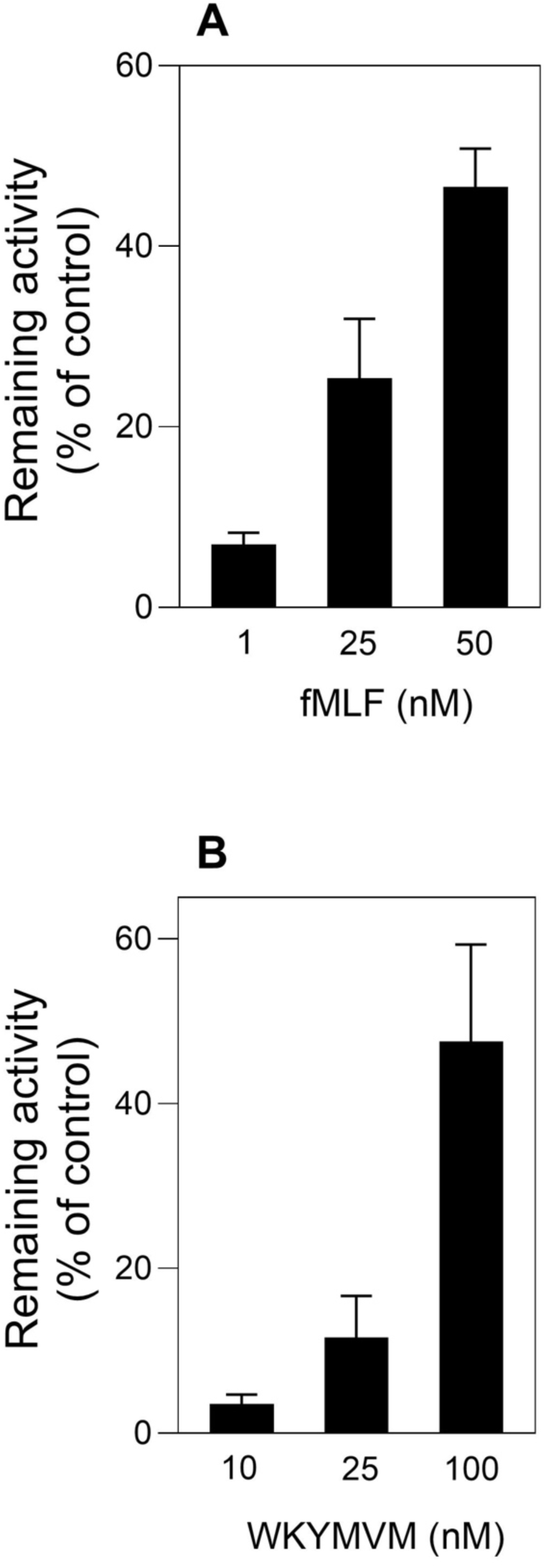
Inhibitory effect of an FPR specific antagonist on the response induced by an FPR agonist combined with the allosteric FFA2R modulator Cmp58. **(A)** Inhibition by the FPR1-specific antagonist cyclosporin H on the response in neutrophils incubated with Cmp58 and induced by different concentrations of fMLF. Inhibition is expressed as the remaining activity (peak response, % of control) after activation with fMLF in the absence and presence of the antagonist (mean ± SEM; n>5). **(B)** Inhibition by the FPR2-specific antagonist PBP10 on the response in neutrophils incubated with Cmp58 and induced by different concentrations of WKYMVM. Inhibition is expressed as the remaining activity (peak response, % of control) of the neutrophil response after activation with WKYMVM in the absence and presence of the antagonist (mean ± SEM; n=3).

The concentration dependent inhibitory profile of the antagonists cyclosporin H (Fig 6A) and PBP10 (Fig 6B) was changed when Cmp58 was introduced in the measuring system (Fig 6).

**Figure 6.**
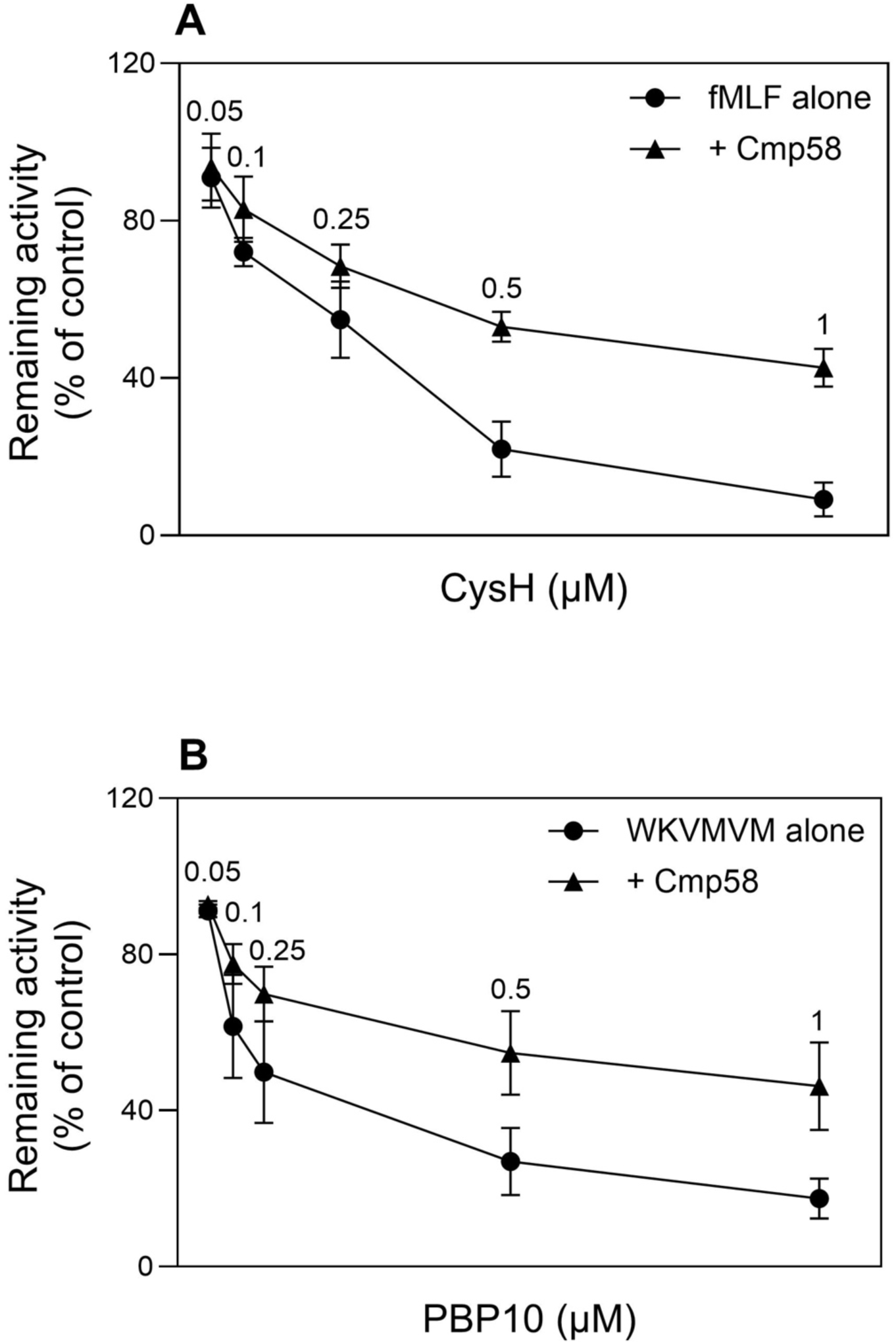
Effect of the allosteric modulator Cmp58 on the inhibition by different concentrations of FPR antagonists when induced by FPR specific agonists. **(A)** Inhibition by different concentrations of the FPR1 specific antagonist cyclosporin H on the neutrophil response induced by fMLF (50nM) in the absence (circles) or presence of Cmp58 (1µM; triangles). Inhibition is expressed as the remaining activity (peak response) in the presence of Cmp58, expressed as percentage of the activity measured in the absence of Cmp58 (mean ± SEM; n=4). **(B)** Inhibition by different concentrations of the FPR2 specific antagonist PBP10 on the neutrophil response induced by WKYMVM (100nM) in the absence (circles) or presence of Cmp58 (1µM; triangles). Inhibition is expressed as the remaining activity (peak response) in the presence of Cmp58, expressed as a percentage of the activity measured in the absence of Cmp58 (mean ± SEM; n>3).

### The FFA2R antagonist CATPB increases the inhibitory effect of the FPR specific antagonists cyclosporin H and PBP10

The lack of a positive effect of Cmp58 response induced by high concentrations of the FPR suggests that FFA2R has no direct effect on this response, but still, its presence changes the inhibitory profile of the FPR antagonists. To determine the involvement of FFA2R in this sensitivity shift, the FFA2R antagonist CATPB was included in the measuring systems to inhibit the effect of the allosteric modulator Cmp58. In the presence of CATPB, the effect of the two FPR antagonists cyclosporin H (Fig 7A) and PBP10 (Fig 7B) returned to the level of inhibition in the absence of Cmp58 (Fig 7).

**Figure 7.**
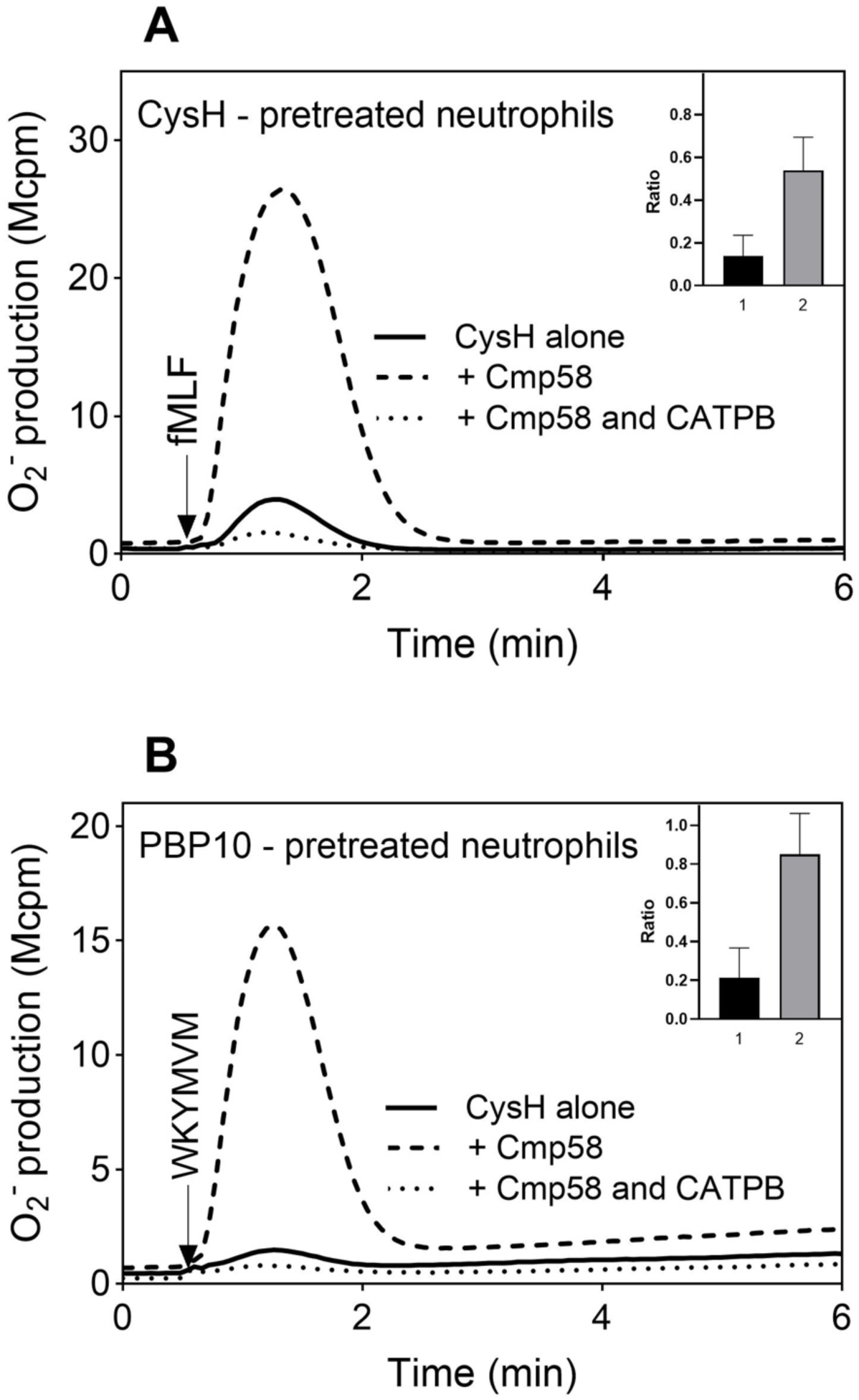
Effect of the FFA2R specific antagonist CATPB on the inhibition pattern of the FPR specific antagonists in neutrophils activated by the corresponding FPR agonist. **(A)** The neutrophil response induced by the FPR1 agonist fMLF (50nM; added at the arrow) in the presence of the FPR1 antagonist (CysH; 1µM) alone (solid line), in combination with the allosteric modulator Cmp58 (1µM; broken line), or together with Cmp58 and the FFA2R- specific antagonist CATPB (100 nM; dotted line), respectively. **Inset:** The inhibition by cyclosporin H (1µM) on the fMLF-induced response is expressed as the ratio between the peak responses obtained in the presence of Cmp58 + CATPB and Cmp58 alone (Ratio 1), and the response obtained in the presence of Cmp58 + CATPB and cyclosporin alone (Ratio 2), respectively. **(B)** The neutrophil response induced by the FPR2 agonist WKYMVM (100 nM; added at the arrow) in the presence of the FPR2 antagonist PBP10 alone (solid line), in combination with the allosteric modulator Cmp58 (1 µM; broken line), or together with Cmp58 and the FFA2R-specific antagonist CATPB (100 nM; dotted line). **Inset:** The inhibition by PBP10 (1µM) on the WKYMVM induced response is expressed as the ratio between the peak responses obtained in the presence of Cmp58 + CATPB and Cmp58 alone (Ratio 1), and the responses obtained in the presence of Cmp58 + CATPB and PBP10 alone (Ratio 2), respectively.

## Discussion

The neutrophil activating peptides fMLF and WKYMVM, recognized by Gα_i_ coupled neutrophil GPCRs, are agonists that when bound to their respective receptor initiate an activation of the superoxide anion producing NADPH oxidase (3). The exact signals generated downstream of the activated receptors FPR1 (receptor for fMLF) and FPR2 (receptor for WKYMVM), that regulate the NADPH oxidase activation upon peptide binding to the receptors have not been fully clarified. It is, however, clear that the receptor downstream induced rise in the cytosolic concentration of free calcium ions ([Ca^2+^]_i_) is not directly involved in this activation. It is also well established that neutrophils are turned into a preactivated state by inflammatory mediators such as bacterial lipopolysaccharide (LPS) and tumor necrosis factor (TNF), and when activated by FPR agonists these primed neutrophils are hyperactive (24). In this study we show that the direct activation of the NADPH oxidase mediated by the FPR specific agonists is not the only mechanisms by which the receptor downstream signals can activate the NADPH oxidase. In the presence of Cmp58, a positive allosteric modulator of FFA2R, the response induced by low-/non-activating concentrations of either of the two peptides is primed. The Cmp58-amplified response induced by the FPR agonist depends both on the respective FPR and FFA2R. This was shown by the inhibition mediated by the FPR1 and FPR2 specific antagonists cyclosporin H and PBP10 as well as by the inhibition mediated by the FFA2R specific antagonist CATPB. Together, these findings support a receptor transactivation mechanism in which signals generated downstream of agonist-occupied FPRs activate the allosterically modulated FFA2R, that in turn activates the NADPH oxidase. The data presented are consistent with an earlier described receptor transactivation model in which intracellular signals generated by several other GPCRs activate FFA2R from the cytosolic side of the plasma membrane, an activation mechanism that works when this receptor is allosterically modulated (16–19). The fact that the FPRs together with FFA2R participate in the peptide-induced activation of the neutrophil NADPH oxidase fully support this receptor transactivation mechanism by which the signals generated by the agonist occupied FPRs indirectly activate the NADPH oxidase through signals generated by FFA2R.

The finding that saturating concentrations of FPRs specific agonists directly activate the neutrophil NADPH oxidase, suggest the presence of two NADPH oxidase activating signaling pathways are induced by the FPRs. One pathway is triggered when many receptors are occupied and these signals directly activate the NADPH oxidase; the other signaling pathway requires few occupied FPRs and the NADPH oxidase is indirectly activated by a receptor transactivation mechanism in which the allosterically modulated FFA2Rs participate. The magnitude of the NADPH oxidase activity induced by high (fully activating) concentrations of the FPR agonists is not affected by the positive allosteric FFA2R, and in agreement with this independence of FFA2R, the response induced is not affected by the FFA2R antagonist CATPB. However, although Cmp58 was without effect on the magnitude of the response mediated by the FPR agonists, the response induced in the presence of Cmp58 was only partly inhibited by the respective FPR and FPR2 antagonist. The inhibitory effect of the two FPR antagonists was, however, restored when the effect of Cmp58 was neutralized/blocked by the FFA2R antagonist CATPB. These data clearly show that the signals that activate the allosterically modulated FFA2Rs are generated when a small fraction of the FPRs are occupied by an agonist, and the signals are of importance for NADPH oxidase activity when the number of agonists that bind and activate the FPR is low; this situation is at hand when the agonist concentration is in the low-/non-activation concentration range, but also when the concentration of the agonist is high but the presence of an FPR specific antagonist hinders agonist binding to many/most of the membrane expressed receptor sites. The receptor transactivation of FFA2R shown to be initiated by the fMLF/FPR1 and WKYMVM/FPR2 agonist/receptor complexes expand the number of GPCRs that in earlier published papers have been shown to utilize such a mechanism to activate the NADPH oxidase in neutrophils. These receptors include P2Y2R (the ATP receptor), PAF (the receptor for platelet activating factor), BLT1 (the LTB_4_ receptor), HCA_3_R (hydroxy-carboxylic acid 3 receptor) and C5aR (the receptor for the complement component C5a) (24). Taken together, these results suggest that it is time that the prevailing paradigm of allosteric receptor modulators, stated that PAMs (positive allosteric receptor modulators), solely affect the response by orthosteric agonists acting on the receptor that recognizes/binds the allosteric modulator (12, 25–27), should be abandoned. Despite signaling differences between the GPCRs shown to transactivate FFA2R, they share the common feature that the downstream signaling cascades include an activation the PLC dependent transient rise in [Ca^2+^]_i_(24). It might, thus, be, that this PLC-PIP_2_-IP_3_ regulated pathway is part of the receptor transactivation mechanism. For this, the FFA2R specific PAM should transfer this receptor to a state that is activated when the cytosolic concentration of free Ca^2+^ ([Ca^2+^]_i_) is increased. This receptor transactivation model is supported by previous findings showing that, when FFA2R is allosterically modulated, an increase in [Ca²⁺]_i_ is sufficient to activate the NADPH oxidase. This was demonstrated using two different approaches to increase [Ca²⁺]_i_ (28); activation by ionomycin (a Ca²⁺ specific ionophore) and thapsigargin (an inhibition of the Ca^2+^-transporting ATPase present in the ER). Taken together, the NADPH oxidase response induced by the potent endogenous activators fMLF and WKYMVM, is regulated by Cmp58, an allosteric modulator of FFA2R; the activation potency is increased without any effect on the efficacy of the FPR agonist. Cmp58 also reduces the inhibitory effect of the specific FPR antagonist, an effect reversed by CATPB, an FFA2R selective antagonist. We conclude that the NADPH oxidase can be activated by two distinct FPR-dependent signaling pathways; one direct and one through transactivation of the allosterically modulated FFA2Rs that secondarily generates signals which elicit NADPH oxidase activity. This is in line with the earlier described receptor transactivation (receptor crosstalk) model, which stipulates that FFA2R is activated by receptor downstream signals from multiple neutrophil GPCRs, now including the two neutrophil FPRs.

## Acknowledgement

The authors thank the former Ph.D. students Simon Lind and Zahra Khan for the discussions about the allosteric modulator Cmp58 and the C5aR mediated transactivation, discussions that inspired us to investigate FPR activity. The authors also thank Linda Bergqvist for technical assistance regarding experiments.

## Credit authorship contribution statement

**Claes Dahlgren**: Conceptualization, Methodology, Validation, Writing – Original draft, Supervision.

**Danni Wang**: Investigation, Visualization, Writing – Review and editing

**Huamei Forsman**: Conceptualization, Methodology, Validation, Investigation, Writing – Review and Editing, Supervision, Funding acquisition.

**Lena Björkman:** Conceptualization, Methodology, Validation, Formal analysis, Investigation, Writing – Review and Editing, Visualization, Supervision, Funding acquisition.

## Abbreviations

CATPB: FFA2R antagonist - (S)-3-[2-(3-chlorophenyl) acetamido]-4-[4- (trifluoromethyl)phenyl]butanoic acid;
Cmp58: FFA2R allosteric modulator - ((*S*)-2-(4- chlorophenyl)-3,3-dimethyl-*N*-(5-phenylthiazol-2-yl)butanamide
FFA2R: free fatty acid receptor 2
GPCRs: G protein-coupled receptors
HRP: horseradish peroxidase
KRG: Krebs- Ringer glucose buffer
O_2_^−^: superoxide anion
PAM: positive allosteric modulator
ROS: reactive oxygen species
SCFAs: short chain fatty acids
TNF: tumor necrosis factor

